# The two extinctions of the Carolina parakeet

**DOI:** 10.1101/801142

**Authors:** Kevin R. Burgio, Colin J. Carlson, Alexander L. Bond, Margaret A. Rubega, Morgan W. Tingley

## Abstract

Due to climate change and habitat conversion, estimates of the number of species extinctions over the next century are alarming. Coming up with solutions for conservation will require many different approaches, including exploring the extinction processes of recently extinct species. Given that parrots are the most threatened group of birds, information regarding parrot extinction is especially pressing. While most recent parrot extinctions have been island endemics, the Carolina parakeet (*Conuropsis carolinensis*) had an 18th-century range covering nearly half of the present-day United States, despite which, they went extinct in the 20th century. The major cause of their extinction remains unknown. As a first step to determining what caused their extinction, we used a newly published, extensive dataset of Carolina parakeet observations combined with a Bayesian extinction estimating model to determine the most likely date of their extinction. By considering each of the two subspecies independently, we found that they went extinct ~30 years apart: the western subspecies (*C. c. ludovicianus*) around 1914 and the eastern subspecies (*C. c. carolinensis*) either in the late 1930s or mid-1940s. Had we only considered all observations together, this pattern would have been obscured, missing a major clue to the Carolina parakeet’s extinction. Since the Carolina parakeet was a wide-ranging species that went extinct during a period of rapid agricultural and industrial expansion, conditions that mirror those presently occurring in many parts of the world where parrot diversity is highest, any lessons we can glean from their disappearance may be vital to modern parrot conservation efforts.

## INTRODUCTION

We are entering the early stages of the “Sixth Mass Extinction” (Ceballos et al. 2015), where estimates of the proportion of species to go extinct over the next century are dire (e.g., Urban 2015, Carlson et al. 2017) and species may not be able to adapt quickly enough to respond to climate change (Keogan et al. 2018, Radchuk et al. 2019). This amount of loss will destabilize already compromised ecosystems, while conservation agencies are finding it difficult to plan for the challenges and uncertainty ahead (Armsworth et al. 2015). While our best guesses about how and when species will go extinct in the future are largely based on model predictions, much can be gained by studying the spatial, temporal, and mechanistic processes that led to recent extinctions (e.g., Stanton 2014, Bond et al. 2019). Investigating the past may not only allow us to recover some of the natural history lost with the extinction of these species but may also yield important insights that can inform conservation actions now and in the future.

Parrots are one of the most threatened orders of birds, with ~ 43% of all species listed as near-threatened or worse by the IUCN (Marsden & Royle 2015), and they face many different pressures, including habitat loss and trapping (Snyder et al. 2000). Parrots are “keystone mutualists,” providing many important ecosystem functions (Tella et al. 2015, Blanco et al. 2016). The loss of parrots would affect plant species especially, contributing to ecosystem instability. While a handful of parrot species have gone extinct over the past few centuries, most were endemic to islands (Olah et al. 2016). There is, however, one notable exception – the Carolina parakeet (*Conuropsis carolinensis*), which had a geographic range an order of magnitude larger than the average for all other recently extinct parrot species (Olah et al. 2016). Even though Carolina parakeets were charismatic and were of considerable interest to ornithologists during the 1800s, we know little about their biology or how and when they went extinct.

The Carolina parakeet is one of the four forest-dependent bird species to go extinct within the continental United States since the arrival of Europeans (Pimm & Askins 1995). Before its decline, the Carolina parakeet was widely distributed, with a range stretching from the mid-Atlantic coast to Nebraska, and south to Florida (Burgio et al. 2017). The Carolina parakeet has two subspecies (*C. c. ludovicianus* and *C. c. carolinensis*) but was not studied in detail while extant. Thus, much of the information regarding their natural history is speculative. Accounts of the species show a gradual population decline starting with European colonization (Snyder and Russell 2002). By the early 1800s, Audubon (1831) noted a marked decline in their population numbers and range, although he still considered them relatively common. During their decline, the species’ range receded from east to west toward the Mississippi River (except in Florida), and from north to south, along the Ohio River, seemingly in concert with the expansion of European expansion, destruction of bottomland forests, and the rise of intensive agriculture (Askins 2000).

It is unclear exactly why the Carolina parakeet went extinct. People shot Carolina parakeets for sport, food, feathers, scientific collections, and to protect crops (Snyder and Russell 2002). However, it seems unlikely that overexploitation was a major contributor to their extinction (McKinley 1980, Snyder 2004). Others have suggested other potential causes, such as competition for food with other avian species, habitat loss, competition with introduced European honey bees (*Apis mellifera*) over nesting/roosting sites, disease, and pressure from trapping for the pet trade (McKinley 1980, Brunswig et al. 1983, Pimm and Askins 1995, Snyder & Russell 2002).

Additionally, it is also unclear *when* the Carolina parakeet went extinct. At the turn of the twentieth century, the Carolina parakeet could be found only in Florida, South Carolina, and a few isolated regions west of the Mississippi River (Hasbrouck 1891, Snyder 2004). The last captive Carolina parakeet died in a zoo about 100 years ago in 1918. The last “accepted” sighting of the parakeet occurred in Florida in 1920 (Snyder 2004), but there were reports of the Carolina parakeet into the 1930s and 1940s in both Florida and the Carolinas (Snyder 2004) which were likely legitimate (Wright 2005). Determining exactly when the Carolina parakeet went extinct is the first step in unraveling the mystery of how they went extinct, a question that may help provide valuable information for parrot conservation and for any future “de-extinction” efforts, for which they are considered a good candidate (Seddon et al. 2014). Here, we use a newly published, extensive dataset of Carolina parakeet occurrence (Burgio et al. 2018), paired with recent advances in extinction date estimation (Solow and Beet 2014, Kodikara et al. 2018), to determine when these iconic parakeets went extinct.

## METHODS

### Data collection

We collected and georeferenced locality data from Carolina parakeet specimens found in natural history collections around the world and observations of Carolina parakeets published in the literature from 1564 to 1944 (see Burgio et al. [2018] for the description of the data collection methods and a link to the freely available dataset). We then split our dataset by subspecies. We considered all occurrence points west of the Appalachian crest and west of Alabama to be *C. c. ludovicianus* and points east of the Appalachian crest and east of the state of Mississippi to be *C. c. carolinensis*. These broad geographic delineations are generally accepted as the range limits of the two subspecies (Ridgway 1916, Swenk 1934), and are consistent with the subspecies identifications listed on all 261 labeled museum specimens represented in the dataset for which subspecies was recorded or inferred.

We determined the level of certainty of each observation based on expert opinion from the literature, from Snyder (2004) and 18 articles by McKinley (1960, 1964, 1965,1976, 1977a, 1977b, 1978a, 1978b, 1978c, 1978d, 1979a, 1979b, 1979c, 1979d, 1981, 1985, McKinley & James 1984, McKinley & Hardy 1985). We truncated our analysis at 1800, as observations before 1800 were sporadic due, primarily, to a lack of consistent occurrence records and also because this was the point at which Audubon (1831) noted decreasing numbers of Carolina parakeets. Within this framework, we used the entire dataset and designated physical evidence, such as a specimen, as “1”, while observations considered legitimate by expert opinion, but not interrogable in the present as “2”, and controversial as “3.” When individual years had multiple records, we always used the evidence with the highest certainty (Figure 1a, 1b). While the last "official" sighting of the Carolina Parakeet was in 1920 (Synder 2004), contemporary experts consider the sightings in the Santee swamp area in South Carolina the 1930s to be legitimate (Snyder 2004, Wright 2005), therefore we treated all reported sightings after 1920 as unconfirmed, aside from those sightings in South Carolina.

**Figure 1.**
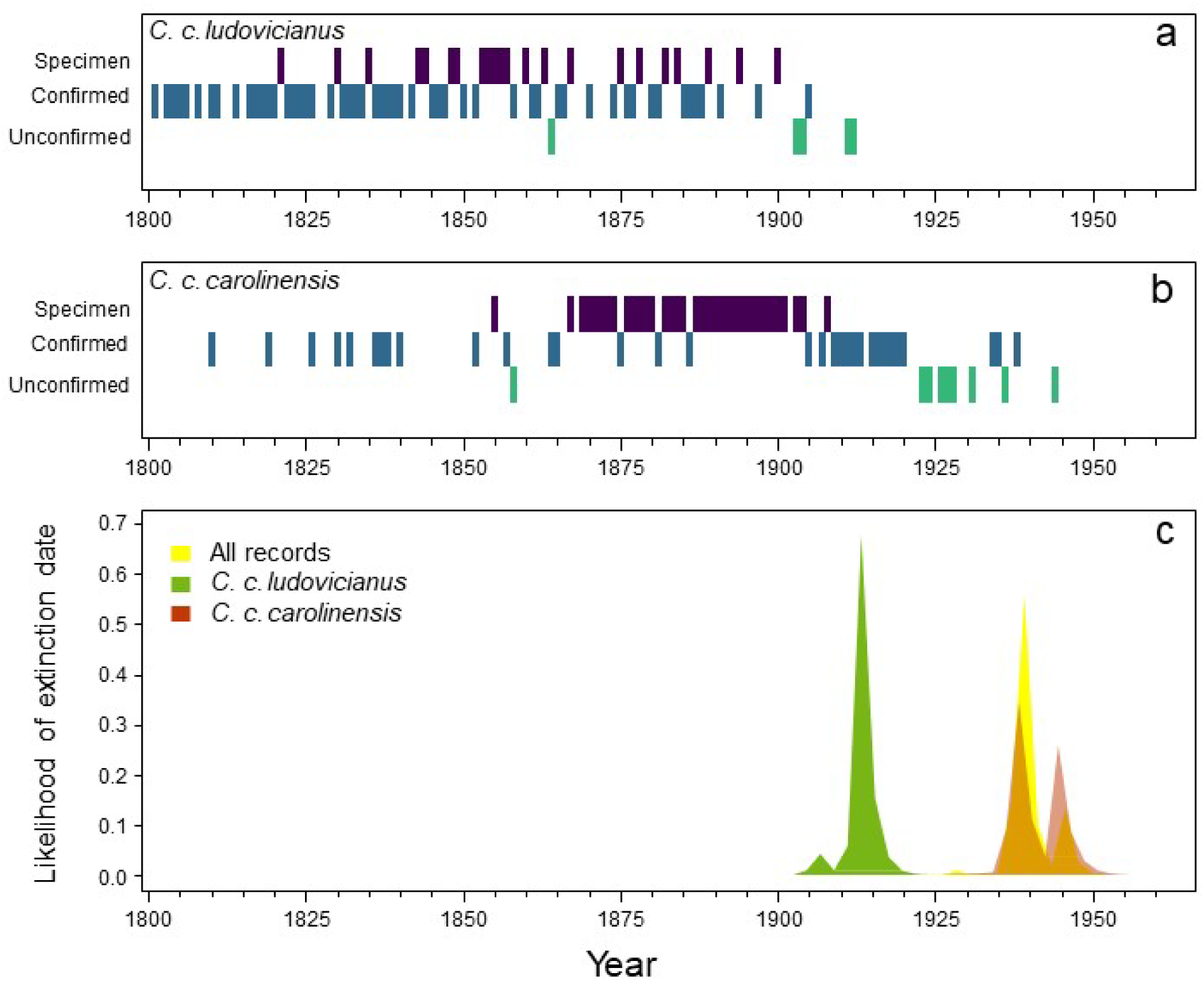
**a.** The sighting record for *C. c. carolinensis*. **b.** The sighting record for *C. c. ludovicianus*. For panels a and b, the top row represents specimen records (purple), the middle row represents confirmed observations (blue), and the bottom row represents unconfirmed or controversial records (green). For each year, we used data with the highest certainty. **c.** The estimates of likely extinction dates for *C. c. ludovicianus* (green), *C. c. carolinensis* (orange), and for all records (yellow).

### Analyses

We estimated the date of extinction for all Carolina parakeets generally and both subspecies specifically. Given that our dataset combined physical evidence (i.e., specimens) and observations with varying degrees of reliability, we used the Bayesian extinction estimating equations proposed by Solow et al. (2012) and modified by Solow and Beet (2014). We ran the analyses in the R package “spatExtinct” (Carlson et al. 2018a). This modeling approach weighs various types of evidence based on their reliability (Solow et al. 2012), and considers the validity of certain and uncertain sightings independently (Boakes et al. 2015, Kodikara et al. 2018), which is especially useful in inferring extinction dates (τ_E_) of recently extinct species with both specimen and observation data (Carlson et al. 2018b, Bond et al. 2019). Of the two models presented in Solow and Beet (2014), we used “Model 2” because some of “uncertain” observations are from reportedly dubious sources (e.g. egg hunters with a vested interest in selling more expensive Carolina parakeet eggs) and “Model 2” is appropriate when uncertain sightings are more likely invalid (Kodikara et al. 2018). We ran these analyses independently for each subspecies (*C. c. carolinensis:* n = 76; *C. c. ludovicianus*: n = 80), and both subspecies together (n = 116).

## RESULTS

We estimated that the eastern subspecies, *C. c. carolinensis*, likely went extinct somewhere in the late 1930s or the mid-1940s, with the two most likely values for τ_E_ in 1938 and 1944 (Fig. 1c) while the western subspecies, *C. c. ludovicianus* went extinct 25-30 years earlier; the most likely value for τ_E_ was 1913 (Fig. 1c). When considering both subspecies together, the extinction estimate doesn’t differ much from the estimate for *C. c. carolinensis* since the highest probability of τ_E_ was 1939 (Fig. 1c); however, failing to consider each subspecies individually obscures important distinctions between the two, especially for inferring causation.

## DISCUSSION

The two separate subspecies of the Carolina parakeet appeared to go extinct ~30 years apart, far later than currently believed, beyond the currently accepted date of 1920 and the most recent analysis that estimated they went extinct around 1915 (Elphick et al. 2010). Elphick et al. (2010), however, used a less complete dataset and did not take into account uncertain sightings. While largely dismissed at the time (Snyder 2004), our results lend credibility to the sightings in South Carolina in the late 1930s. It is even possible that the observation from North Carolina in 1944 reported to Roger Tory Peterson (Snyder 2004) may have been accurate, as well as the mystery population of Carolina parakeets in Florida reported by ornithologist Oscar Baynard to persist well into the late 1930s, but for which he refused to disclose the location (Snyder 2004). That the western subspecies went extinct first despite occupying a larger range (Burgio et al. 2017) is curious, and suggests a lower initial population, or different or more severe pressures.

The Carolina parakeet continues to be one of the most popular proposed targets for de-extinction projects (Seddon et al. 2014). Aside from the ethical issues surrounding de-extinction generally (see Blockstein 2017), unless researchers can identify the major driver of their extinction, de-extinction efforts for the Carolina parakeet may be doomed to fail at great expense in both time and money. Despite a lack of evidence, many believe that farmers, egg poachers, and trappers pushed the species to extinction (McKinley 1980, Snyder 2004), buoyed in part by the lurid anecdote of a farmer shooting the parakeets in his orchard found in Audubon’s foundational book on North American birds (Audubon 1831). While the evidence is indeed scant, researchers who have focused on this species over the past 60+ years point to a disease likely transmitted from domestic poultry as the most likely cause but have not settled on which disease (McKinley 1980, Snyder and Russell 2002). Finally, the landscape across the parakeet’s former range has shifted dramatically since their decline started around 1800, which would make de-extinction efforts difficult at best, especially when considering the uncertainty associated with climate change.

Learning the most likely extinction dates of the two subspecies of the Carolina parakeet is the first of many steps needed to solve the mystery that has eluded researchers for well over 50 years. That the two subspecies went extinct 30 years apart is an important clue which has implications for how we interpret the loss of this species and consider the factors that pushed a species that appeared stable in 1800 to be found only in small populations in remote areas 100 years later. The limitations in both landscape change data and occurrence data may make it difficult to rule out habitat loss as a major factor (but see Pimm and Askins 1995). Advances in genetic analyses, as seen recently seen with passenger pigeons (*Ectopistes migratorius*; Hung et al. 2015), or examining the few preserved specimens in natural history museums for signs of disease (e.g., Rothschild & Panza 2005) may help shed light on what ultimately drove the Carolina parakeet to extinction. The Carolina parakeet was a wide-ranging species and faced challenges of the rapid agricultural expansion and industrialization of the United States during the 20th century, a process being repeated today in many areas where parrots are found. Given that parrots are so threatened, determining the cause of the Carolina parakeet extinction is especially pressing.

## ACKNOWLEDGMENTS

K. Burgio was supported by a National Science Foundation GRFP grant # DGE-0753455 and a National Science Foundation NRT-IGE grant #1545458 to M. Rubega.

